# Multi-Modal Large Language Model Enables Protein Function Prediction

**DOI:** 10.1101/2024.08.19.608729

**Authors:** Mingjia Huo, Han Guo, Xingyi Cheng, Digvijay Singh, Hamidreza Rahmani, Shen Li, Philipp Gerlof, Trey Ideker, Danielle A. Grotjahn, Elizabeth Villa, Le Song, Pengtao Xie

## Abstract

Predicting the functions of proteins can greatly accelerate biological discovery and applications, where deep learning methods have recently shown great potential. However, these methods predominantly predict protein functions as discrete categories, which fails to capture the nuanced and complex nature of protein functions. Furthermore, existing methods require the development of separate models for each prediction task, a process that can be both resource-heavy and time-consuming. Here, we present ProteinChat, a versatile, multi-modal large language model that takes a protein’s amino acid sequence as input and generates comprehensive narratives describing its function. ProteinChat is trained using over 1,500,000 (protein, prompt, answer) triplets curated from the Swiss-Prot dataset, covering diverse functions. This novel model can universally predict a wide range of protein functions, all within a single, unified framework. Furthermore, ProteinChat supports interactive dialogues with human users, allowing for iterative refinement of predictions and deeper exploration of protein functions. Our experimental results, evaluated through both human expert assessment and automated metrics, demonstrate that ProteinChat outperforms general-purpose LLMs like GPT-4, one of the flagship LLMs, by over ten-fold. In addition, ProteinChat exceeds or matches the performance of task-specific prediction models.

## Introduction

Proteins, composed of amino acid sequences that determine their unique structures and functions, are fundamental molecules essential for life-sustaining processes. Understanding protein functions and properties (collectively referred to as functions in this manuscript for simplicity) is crucial for advancing biological knowledge and driving innovations in drug discovery, disease treatment, and synthetic biology (1–5). Predicting protein functions is a complex and challenging task due to the inherent diversity and intricate nature of proteins (6–10). Recent advancements in deep learning have demonstrated significant potential in improving the accuracy and efficiency of protein function prediction (11–18). By leveraging extensive datasets of protein sequences, structures, and annotated functions, deep learning models can discern intricate patterns and relationships that often elude traditional computational methods. The success of tools like CLEAN (17), which predicts enzyme functions with superior accuracy compared to traditional methods like BLASTp (19), exemplifies the transformative impact of deep learning in the field.

However, existing deep learning-based methods for protein function prediction face significant limitations that prevent them from fully capturing the diverse range of protein functions. These methods typically predict protein functions as discrete categories (7, 12, 13, 16–18). This oversimplification fails to reflect the complex and nuanced nature of proteins which often perform multiple functions, engage in various interactions, and participate in intricate biological pathways. Additionally, existing methods necessitate the development of specialized models for each prediction task, resulting in a fragmented approach that lacks efficiency and scalability (8, 13, 15–18). The absence of a unified model capable of concurrently handling various prediction tasks limits a holistic understanding of protein functions. This fragmentation also increases the complexity and resource requirements for research and development, as developing, training, and maintaining multiple specialized models is significantly more challenging than managing a single, versatile model.

Large language models (LLMs) (20–22) hold significant potential for addressing the limitations of current deep learning-based protein function prediction methods. These LLM models excel in generating high-quality text, making them well-suited for describing complex protein functions through comprehensive narratives. Furthermore, a single, pretrained LLM can perform a wide array of prediction tasks using task-specific user instructions or questions described in natural language (referred to as *prompts*) (23, 24), eliminating the necessity of training separate models for each task. Furthermore, LLMs facilitate interactive dialogues with human users (25, 26), enabling iterative refinement of generated textual predictions.

We developed ProteinChat, a multi-modal LLM that integrates two modalities - protein sequences and text. It takes an amino acid sequence and a prompt as inputs, and generates a detailed textual prediction of the protein’s function. Unlike traditional methods that predict protein functions as discrete categories, ProteinChat generates coherent and comprehensive texts to predict the multi-faceted functions of proteins, capturing the detailed roles, interactions, and biological context of proteins in a manner akin to human expert descriptions. Moreover, ProteinChat enables the use of diverse prompts for various prediction tasks that cover a wide range of protein functions and properties within this single tool, thereby streamlining the whole protein function exploration process without requiring new model training or extensive maintenance. Significantly outperforming current methods including GPT-4 (24), ProteinChat can make accurate predictions across a broad spectrum of protein functions, which were evaluated using multiple metrics including assessments by human experts.

## Results

### ProteinChat overview

ProteinChat accepts two types of inputs simultaneously: the amino acid sequence of a protein and a prompt tailored for easy, human-like dialogues with ProteinChat. For example, when given the prompt “describe the functions of this protein”, ProteinChat generates a detailed free-form text describing the protein’s various functions (Fig. 1a). Besides free-form prediction, ProteinChat can also predict specific function categories. For example when prompted with “What type of enzyme is this? Choose from [*a list of categories*]”, ProteinChat chooses a specific answer from the list (Fig. 1a).

**Fig. 1.**
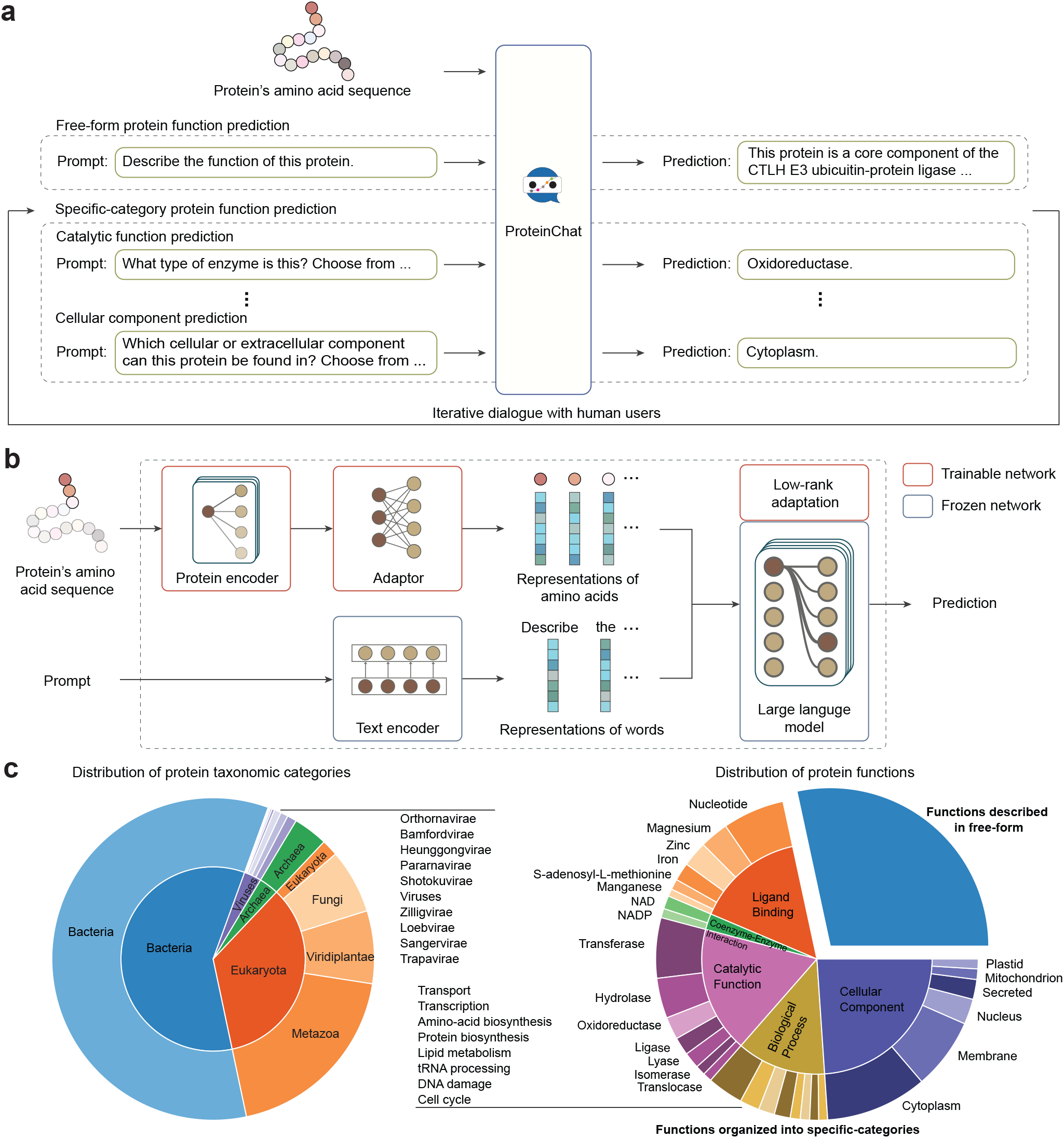
ProteinChat is a multi-modal LLM capable of predicting protein functions represented either in free-form text or as specific categories. **a**, ProteinChat enables versatile prediction of protein functions, allowing users to submit various requests in flexible natural language (known as prompts). By using task-specific prompts, ProteinChat can perform a variety of prediction tasks within a single framework without changing model parameters. ProteinChat facilitates interactive dialogues with users by retaining the conversation history, including prompts and corresponding predictions, allowing for in-depth analysis of a specific protein over multiple interactions. **b**, Model architecture of ProteinChat. It takes the amino acid sequence of a protein and a prompt as inputs, then generates a prediction in natural language. ProteinChat consists of a protein encoder that learns representation vectors for amino acids (AAs), an adaptor that transforms these representations into a format compatible with LLMs, and an LLM that generates the prediction based on the AAs’ representations and the prompt. **c**, An extensive dataset, comprising proteins from various taxonomic groups, was constructed to train ProteinChat. In the left pie chart, the inner ring represents superkingdoms, while the outer ring represents kingdoms. ProteinChat was trained to make two types of predictions: one generates free-form textual descriptions, and the other predicts specific function categories. The pie chart on the right displays the relative proportions of the training data devoted to these two types.

ProteinChat consists of three key modules: a protein encoder, an LLM, and an adaptor that bridges the two (Fig. 1b). The protein encoder processes the amino acid sequence of the input protein, generating a representation vector for each amino acid, which captures the molecular characteristics of that amino acid. The adaptor aligns these representations with the LLM by transforming them into a format that is compatible with the LLM’s input. Once this alignment is achieved, the LLM integrates the amino acid sequence with the prompt, and then utilizes this combined input to generate a textual prediction of the protein’s function. We utilized xTrimoPGLM (27), a state-of-the-art protein language model, as the protein encoder, and Vicuna-13B (25), fine-tuned from Llama-2 (21), as the LLM of ProteinChat.

To train the ProteinChat model, we assembled a comprehensive dataset comprising (protein, prompt, answer) tripletts sourced from the Swiss-Prot database (28), the expertly curated section of UniProt Knowledgebase (UniProtKB) (29). The dataset contains approximately 1.5 million triplets from 523,994 proteins. In each triplet, the protein and prompt serve as inputs to the ProteinChat model, while the answer represents the desired output of ProteinChat. The answer can be either a detailed free-form text describing protein functions or a UniProtKB keyword representing a specific function category. This dataset comprehensively encompasses a diverse taxonomy of proteins and their various functions (Fig. 1c).

For the pretrained LLM (Vicuna-13B), we applied Low-Rank Adaptation (LoRA) (30) for fine-tuning. Specifically, a low-rank update matrix was added to each pretrained weight matrix. During fine-tuning, only the low-rank matrix was updated, while the original pretrained weight matrices remain fixed. For the pretrained protein encoder (xTrimoPGLM), full fine-tuning was utilized: all the pretrained weights were updated. The adaptor was trained from scratch. The trainable weights were optimized by minimizing the negative log-likelihood loss between the input data (proteins and prompts) and the corresponding output answers. Further details on the training of ProteinChat are provided in Methods.

### ProteinChat’s free-form predictions vastly outperform GPT-4

Using the prompt “please describe the function of this protein”, ProteinChat generated free-form text predictions for the functions of 200 randomly selected proteins from Swiss-Prot. These proteins were not included in the training data. The random selection process resulted in a diverse set of proteins with a wide range of functions. The generated textual predictions offer more specific details about protein functions compared to discrete categories like Enzyme Commission (EC) numbers (17) and Gene Ontology terms (31, 32). As mentioned before, Swiss-Prot includes a textual description of each protein’s function, which was used as ground truth in our evaluation. For a comparative analysis between ProteinChat and GPT-4 (a flagship LLM), we utilized GPT-4 to predict protein functions using two types of inputs: amino acid sequences as strings and protein names. The prompts used for GPT-4 are provided in Methods. We performed a human assessment of the predictions generated by both ProteinChat and GPT-4, where experts specializing in proteins compared the predictions with the corresponding ground truth. They assigned scores of 2, 1, 0, or *Ambiguous* to each prediction. A score of 2 is given when the prediction completely matches, partially matches, adds accurate details to, or provides a credible alternative to the ground truth. A score of 1 is assigned when the prediction is partially correct but contains inaccuracies compared to the ground truth. A score of 0 is assigned when a prediction is completely inaccurate or irrelevant to the ground truth. The *Ambiguous* score is used when it lacks sufficient information to make a comparison between the prediction and the ground truth. A detailed description of the assessment rubric can be found in Extended Data Table 2. Fig. 2c provides examples illustrating how these scores were assigned.

**Fig. 2.**
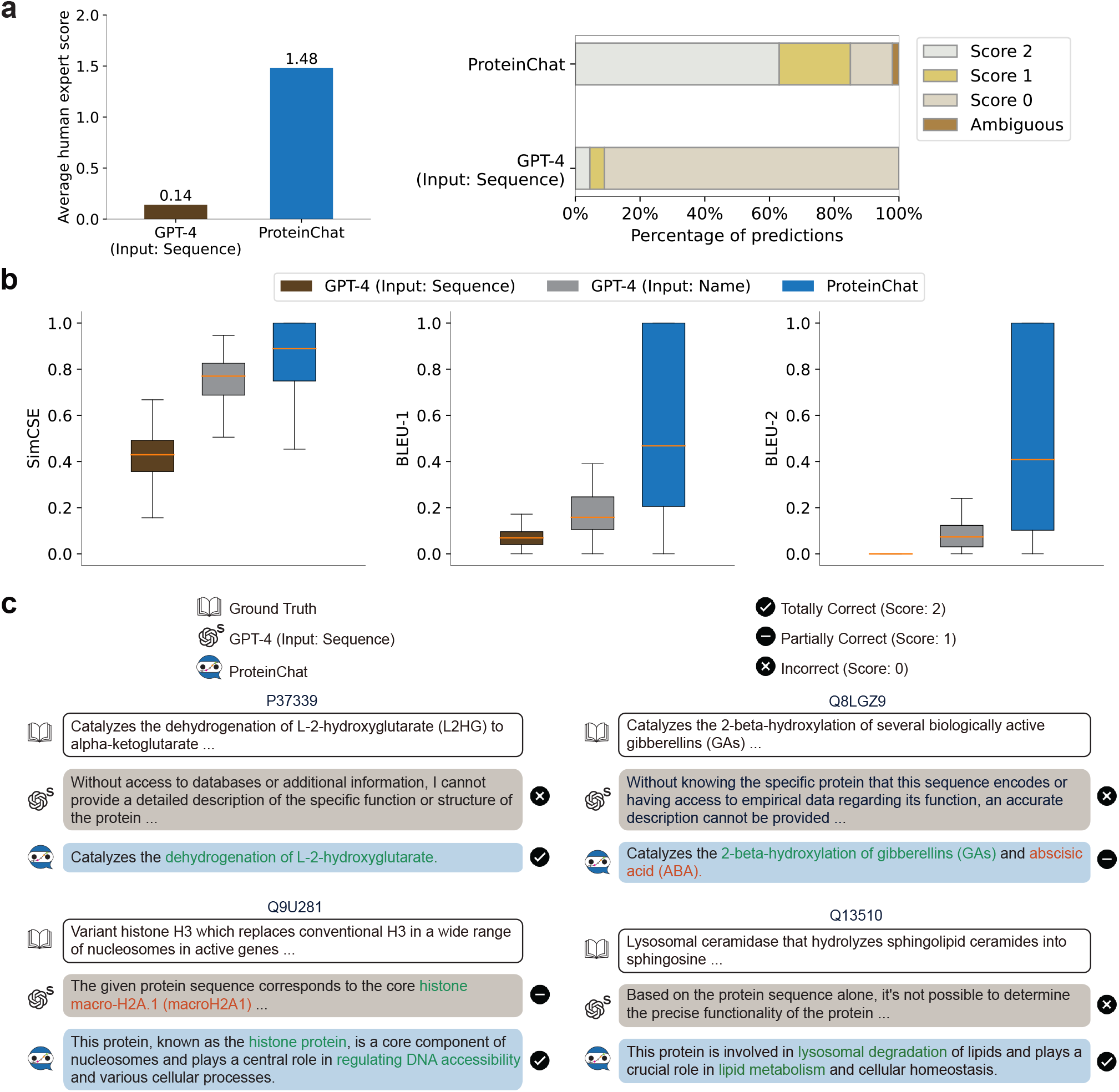
ProteinChat accurately predicts protein functions expressed in textual descriptions and outperforms GPT-4. **a**, ProteinChat significantly outperforms GPT-4 in human expert assessments, by more than ten-fold. Experts assessed predictions on a 0-2 scale: 2 for completely correct, 1 for partially correct, and 0 for incorrect. The average scores are on the left, with the distribution of scores on the right. Like ProteinChat, GPT-4 uses amino acid sequences of proteins as input. **b**, In automated evaluation metrics including SimCSE, BLEU-1, and BLEU-2, ProteinChat demonstrates significantly superior performance compared to GPT-4 which uses amino acid sequences or protein names as inputs. **c**, Examples of predictions generated by ProteinChat and GPT-4 demonstrate that ProteinChat’s predictions are more accurate and informative than those of GPT-4.

ProteinChat achieved an average human assessment score of 1.48, significantly outperforming GPT-4, which had a score of 0.14, by more than ten times. The distribution of scores further highlights the substantial difference between the two models. For ProteinChat, the percentage of proteins that received scores of 2, 1, 0, and Ambiguous were 63%, 22%, 13%, and 2%, respectively. In comparison, GPT-4’s corresponding percentages were 4.5%, 4.5%, 91%, and 0%.

In addition to human assessment, we employed two widely used automated metrics, SimCSE (33) and BLEU (34), to assess the similarity between predicted and ground truth functions for both ProteinChat and GPT-4. SimCSE assesses semantic similarity by comparing the contextual embeddings of texts, generating scores ranging from -1 to 1, with higher values indicating stronger semantic similarity. BLEU, which scores between 0 and 1 with higher values indicating better performance, assesses lexical similarity by comparing n-grams. ProteinChat achieved average SimCSE, BLEU-1, and BLEU-2 scores of 0.85, 0.55, and 0.51 respectively, substantially outperforming GPT-4, which scored 0.42, 0.07, and 0.01 with protein sequences as input, and 0.74, 0.18, and 0.08 with protein names as input (Fig. 2b).

Fig. 2c and Extended Data Fig. 5 present the predictions made by ProteinChat and GPT-4 for some randomly selected proteins, with human expert assessments. These proteins have widely distinct functions and properties. ProteinChat’s predictions consistently surpass those of GPT-4 for these proteins. Specifically, the predictions made by GPT-4 were significantly non-specific, uninformative, and inaccurate. For example, it responded with statements like, “without access to databases or additional information, I cannot provide a detailed description of the specific function or structure of the protein”. In contrast, the predictions made by ProteinChat accurately describe protein functions with rich detail and specificity, closely aligning with the ground truth. For example, ProteinChat’s prediction for protein P37339 received a human assessment score of 2 (“totally correct”). ProteinChat accurately identified the protein’s catalytic functions and specified that its catalytic activity involves the dehydrogenation of L-2-hydroxyglutarate, which aligns very well with the ground truth. In contrast, the response from GPT-4 is uninformative. ProteinChat’s prediction for protein Q8LGZ9 received a score of 1 (“partially correct”): it accurately predicted that the protein catalyzes the 2-beta-hydroxylation of gibberellins (GAs); however, it incorrectly predicted that the protein also catalyzes the 2-beta-hydroxylation of abscisic acid (ABA). Despite this error, the prediction is still significantly more informative than that of GPT-4, which provided no useful insights. Notably, among ProteinChat’s predictions scored as 1, 86% accurately identified the core function but lacked precision on specific details. For example, ProteinChat correctly identified the function or reaction but misattributed the substrate or location, or pinpointed the biological process but failed to specify the involved protein.

Furthermore, the predictions in Fig. 2c illustrate that, unlike previous methods that predict protein functions as discrete categories, ProteinChat generates cohesive and thorough natural language narratives about the diverse functions of proteins. Previous methods often fall short in capturing the complexity and nuance of protein functions, as they reduce these functions to simplistic categories. ProteinChat, however, generates rich, detailed descriptions that mirror the comprehensive analyses provided by human experts. This capability allows for a more holistic understanding of proteins, encompassing their intricate roles, interactions, and biological significance. By utilizing large language models, ProteinChat describes the multifaceted nature of proteins in a way that is both accessible and scientifically rigorous. This method enhances our understanding of individual proteins and facilitates insights into the broader biological systems they operate within. The results from human expert assessments, automated evaluations, and qualitative examples all clearly demonstrate that ProteinChat significantly outperforms GPT-4. This superior performance is primarily due to ProteinChat’s enhanced ability in interpreting a fundamental language of biology, i.e., protein sequences (translated from DNA sequences). As a multi-modal LLM, ProteinChat is specifically designed to understand the amino acid sequences of proteins through a specialized Protein Language Model (PLM) and articulates its understanding via a comprehensive LLM. The PLM is specifically trained on vast datasets of protein sequences, allowing it to capture intricate biochemical relationships and patterns that are essential for accurate protein function prediction. This specialized training enables ProteinChat to offer precise annotations, identify functional domains, and predict potential interactions with high accuracy. Additionally, ProteinChat’s ability to integrate and synthesize data from various sources, including structural databases and functional annotations, further enhances its predictive capabilities. In contrast, GPT-4 treats amino acid sequences merely as strings of letters, relying on a general textual language model for interpretation, which results in a markedly inferior ability in comprehending proteins. Despite its impressive linguistic prowess, GPT-4 lacks the domain-specific training and the multi-modal capabilities that ProteinChat possesses. GPT-4’s general text-based approach to interpreting amino acid sequences means it can miss subtle but crucial biochemical nuances, leading to less reliable predictions. Although GPT-4’s predictions based on protein names were more informative, they are still less specific than those of ProteinChat. It is worth noting that protein names often reveal real protein functions, giving GPT-4 an unfair advantage compared to ProteinChat. In theory, GPT-4 (using protein name) can only work for well-known proteins with extensive, well-documented literature, which was presumably used to train GPT-4. It cannot respond well to novel or undocumented proteins, as there was no prior literature to feed its training. These novel proteins are the bedrock of future scientific discoveries, thus marking a significant limitation of general-purpose LLMs in driving innovation in proteomics. In contrast, ProteinChat is built upon amino acid sequences, a more fundamental feature of proteins, enabling it to understand novel proteins and predict their functions accurately. We also utilized other metrics to evaluate ProteinChat, including assessments by GPT-4 (Extended Data Fig. 2a) and biological term accuracy (Extended Data Fig. 2b), where ProteinChat demonstrated superior performance. Visualizations (Extended Data Fig. 3) demonstrate that ProteinChat effectively groups functionally similar proteins together in its protein representation space, facilitating the accurate prediction of protein functions.

### ProteinChat excels in predicting discrete function categories with high accuracy

In some databases, certain protein functions are organized into discrete categories. For example, in UniProtKB, the catalytic functions of enzymes are categorized as hydrolases, oxidoreductases, lyases, and others. While ProteinChat is designed as a general-purpose tool for generating detailed and nuanced descriptions of a protein’s functions, it can also be customized for specific protein function prediction tasks where functions are categorized discretely. This can be achieved by appropriately adjusting the prompts. We applied ProteinChat to five specific protein function/property prediction tasks curated from UniProtKB, including catalytic function prediction, ligand binding function prediction, coenzyme-enzyme interaction prediction, biological process prediction, and cellular component compartmentalization prediction. These tasks encompass a broad spectrum of protein functions/properties (Methods). It is important to note that these prediction tasks are not mutually exclusive and can overlap. For instance, a particular catalytic function might involve specific ligand binding, or a catalytic function could be a part of a broader biological process.

To accomplish these tasks, we designed task-specific prompts (Methods) for ProteinChat, following a similar style. For enzyme catalytic function prediction, the prompt is “What type of enzyme is this? Choose from [*a list of categories*]”. For biological process prediction, the prompt was: “What biological process is this protein involved in? Choose from [*a list of categories*]”. ProteinChat then selects a specific answer from the given list of categories. The discrete nature of these categories allowed us to objectively evaluate ProteinChat’s performance in comparison to other methods. We employed accuracy, macro F1 score, and weighted F1 score as evaluation metrics, with F1 scores specifically accounting for both false positives and false negatives. We also developed specialized classifier models, each designed to perform a specific prediction task, to evaluate how well ProteinChat, as a more general-purpose model, compares to these task-specific models.

Across all five prediction tasks, ProteinChat demonstrated near-optimal performance (Fig. 3a). It achieved accuracy, macro F1, and weighted F1 scores within the range of 0.95 to 0.99. In contrast, GPT-4’s performance was significantly lower when provided with either a protein name or an amino acid sequence as input. Additionally, ProteinChat either outperformed or matched the results of specialized classifiers, which is particularly remarkable given that ProteinChat employs a single model to handle all these prediction tasks, whereas the specialized classifiers are individually trained for each different task. Developing a specialized model for each prediction task involves extensive training data collection, model training, and hyperparameter tuning, which is time-consuming, resource-intensive, and requires significant domain expertise to ensure accuracy and reliability. Additionally, specialized models cannot easily adapt to new or related tasks without undergoing the entire development process again. In contrast, ProteinChat leverages a single model to perform a variety of protein function prediction tasks by simply modifying the prompts, thereby eliminating the need for developing separate models for each task. This enhances efficiency, flexibility, and scalability.

**Fig. 3.**
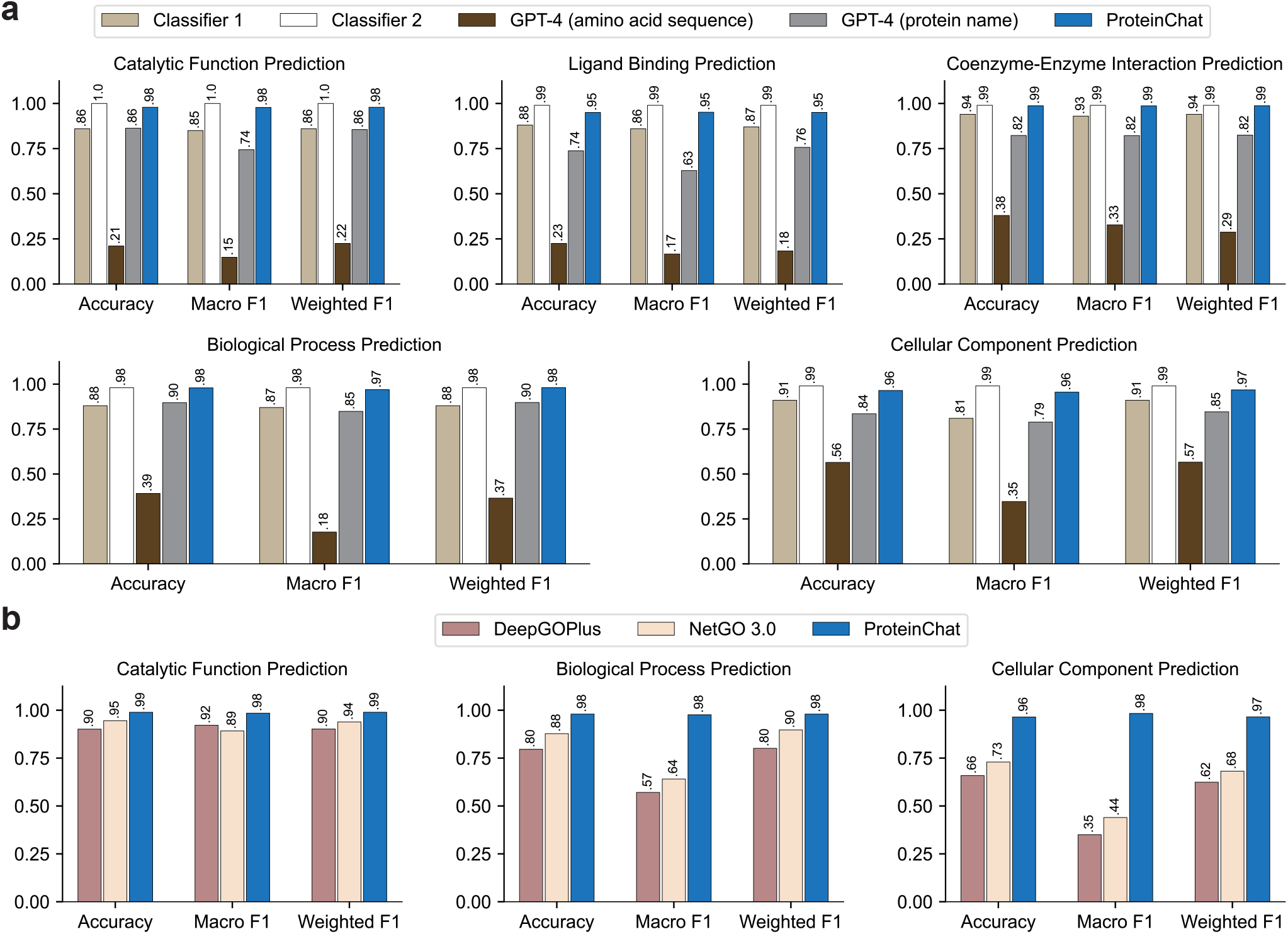
ProteinChat demonstrates exceptional accuracy in specific-category predictions, significantly outperforming GPT-4 and specialized classifiers. **a**, In five specific prediction tasks curated from UniProt, including catalytic function prediction, ligand binding prediction, coenzyme-enzyme interaction prediction, biological process prediction, and cellular component prediction, where protein functions are represented as discrete categories, ProteinChat achieves significantly better accuracy, macro F1, and weighted F1 scores compared to GPT-4 and specialized classifiers. **b**, In predicting protein functions represented using Gene Ontology (GO) categories, ProteinChat significantly outperforms two state-of-the-art GO classifiers - DeepGOPlus and NetGO 3.0.

Next, we utilized ProteinChat to predict protein functions/properties represented by discrete Gene Ontology (GO) (31) categories and compared its performance against leading GO classifiers, including DeepGOPlus (35) and NetGO 3.0 (36). Gene Ontology (GO) is a database that provides a hierarchical structure of categories widely used for annotating protein functions/properties. ProteinChat significantly outperforms DeepGOPlus and NetGO 3.0 in predicting catalytic functions, biological processes, and cellular components (Fig. 3b). For example, ProteinChat achieves a macro F1 score of 0.98 in predicting biological processes, significantly outperforming DeepGOPlus and NetGO, which have scores of 0.57 and 0.64, respectively. ProteinChat outperforms both DeepGOPlus and NetGO due to its ability in retaining and processing the entire sequence of amino acid representations using a protein language model. This ability allows ProteinChat to capture intricate relationships, positional context, and long-range dependencies within the sequence, which are essential for accurate protein function/properties prediction. In contrast, NetGO 3.0 averages the representations into a single vector, losing important sequence information and contextual relationships. DeepGOPlus utilizes convolutional neural networks (CNNs) to learn representations for amino acids, which falls short in capturing long-range dependency between amino acids when compared to the Transformer (37) based protein encoder employed in ProteinChat.

### ProteinChat enables interactive and iterative predictions of protein functions

ProteinChat facilitates interactive dialogues between users and the system. After obtaining the initial predictions from ProteinChat, users can input more detailed and specific prompts to further refine and expand these predictions. Fig. 4 presents three example dialogues between ProteinChat and human users, corresponding to proteins Q9U281, Q9XZG9, and Q9LU44 in UniProtKB. The dialogue on the left pertains to Q9U281, where the user inquires about the general function of this protein. ProteinChat identifies it as a histone protein involved in modulating DNA accessibility. Subsequently, the user inquires about the specific functions of this histone protein, and ProteinChat provides detailed predictions, highlighting the protein’s roles in transcription regulation and post-translational modifications. The top right dialogue pertains to Q9XZG9, where ProteinChat initially predicts that the protein has antibacterial function. Based on the user’s further prompt, ProteinChat then accurately predicts the protein can inhibit the growth of both Gram-positive and Gram-negative bacteria. The bottom right example focuses on Q9LU44. When inquired about general functions, ProteinChat predicts that the protein is involved in pre-mRNA splicing. Upon further inquiry into specific molecular functions, such as metal binding, ProteinChat predicts that the protein binds zinc ions. This dynamic interaction between ProteinChat and users facilitates continuous, in-depth analysis of the same protein, in contrast to previous methods that offer only single-shot predictions. Users can delve deeper into the specifics of protein functions, exploring intricate details and nuances that single-shot predictions might miss. This ensures that the predictions are not only more accurate but also more comprehensive, uncovering complex protein behaviors and mechanisms.

**Fig. 4.**
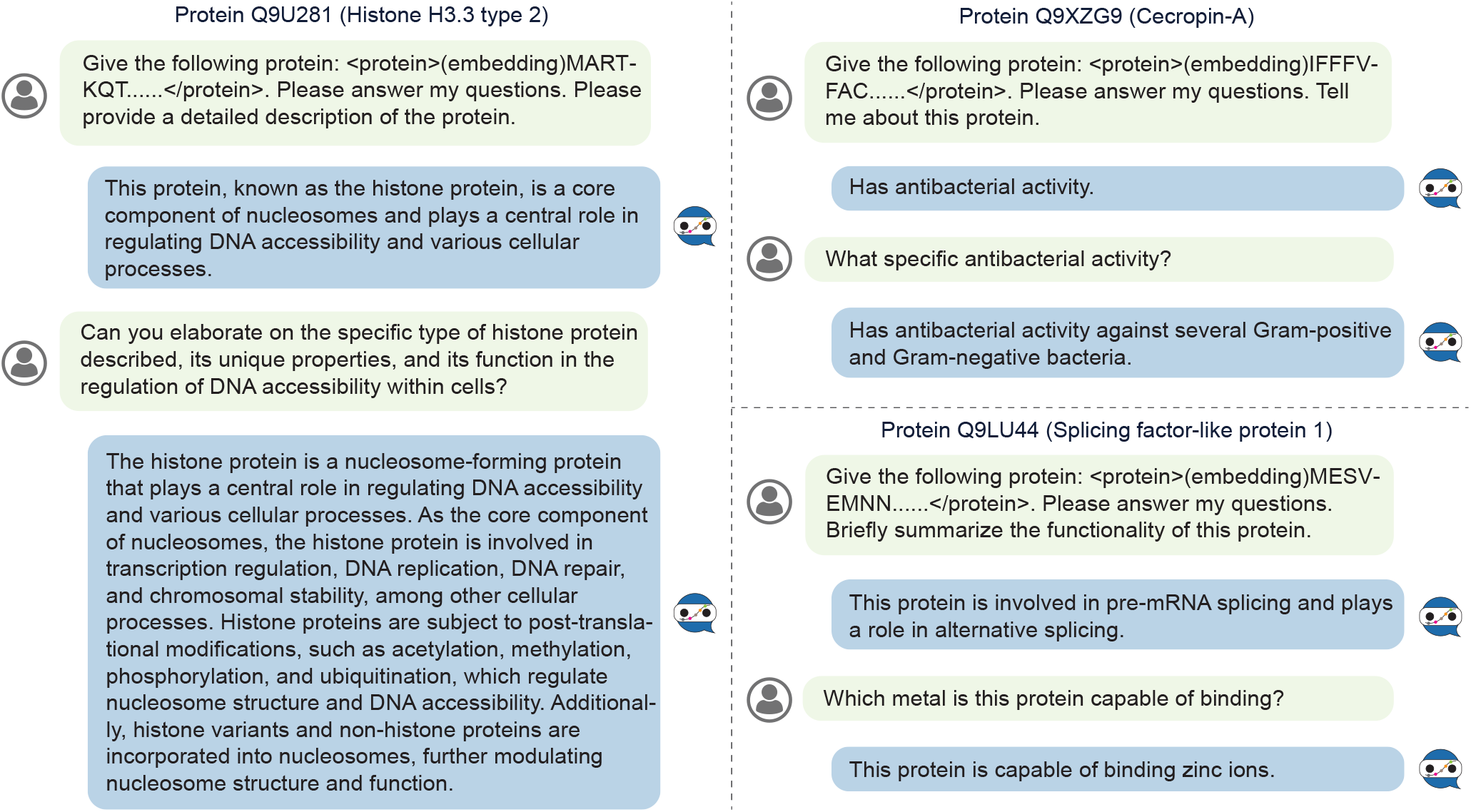
Interactive dialogues between ProteinChat and human users about proteins Q9U281, Q9XZG9, and Q9LU44.

## Discussion

ProteinChat illustrates two important concepts. Firstly, the fundamental language of biology - amino acid sequences-encodes highly rich information about underlying biological processes. This information is both computable and predictive, suggesting that this language can be harnessed to develop powerful predictive models in other areas of biology, as demonstrated by ProteinChat. Secondly, achieving a balance is crucial when designing deep learning models for biological applications. While highly specialized models like DeepGo or NetGo are effective in specific tasks, they may overlook the complex, multi-tasking nature of proteins that are involved in multiple biological pathways. On the other hand, overly generalized models, such as GPT-4, might lack the precision needed for accurate, domain-specific predictions. ProteinChat strikes a balance between these extremes, offering broad generalization across proteomics while maintaining high accuracy and specificity, as demonstrated in Fig. 2 and 3.

ProteinChat is designed to minimize the need for continuous user training while allowing for periodic updates and enhancements by us, the developers. For example, we plan to integrate more advanced versions of Llama (e.g., Llama-3 (38)) as the textual LLM component of ProteinChat, improving the quality of human-like interactions. Additionally, incorporating newer versions of xTrimoPGLM will further enhance ProteinChat’s accuracy and specificity. These planned improvements will ensure that ProteinChat remains both competitive and up-to-date. Furthermore, ProteinChat’s versatility enables seamless integration with other deep-learning models, such as those based on structure prediction like AlphaFold (39), allowing it to predict the functions of proteins in the context of their 3D structures.

Some predictions made by ProteinChat, currently labeled as incorrect by human experts, may actually uncover previously unidentified properties and functions of these proteins. As a result, the scores we assigned to ProteinChat could potentially be even higher. More importantly, predictions deemed incorrect might actually offer new insights or hypotheses that warrant further experimental validation. For many proteins, only a portion of their amino acid sequences have been fully understood, with the remainder still elusive and sometimes labeled as “junk”-sequences that seemingly do not contribute significantly to the protein’s main function (40). ProteinChat has the potential to shed light on these currently uninterpretable sequences. Additionally, large portions of proteins can consist of disordered segments-sequences that do not fold into a stable structure (41). Historically, these segments have often been truncated in structural and biophysical studies, leading to incomplete characterizations. However, recent research indicates that these disordered segments are crucial for the phase separation of proteins into specific cellular compartments, where they carry out their functions (42). ProteinChat, which can analyze the entire protein sequence, could be particularly effective in interpreting these disordered segments and predicting their phase-separating characteristics. This capability may already be reflected in ProteinChat’s predictions related to cellular compartmentalization.

In conclusion, we present ProteinChat, a versatile tool for predicting protein functions represented in text using a multi-modal large language model. ProteinChat provides nuanced and in-depth predictions, surpassing both general-purpose LLMs and task-specific classifiers. Its ability in handling various prediction tasks within a single framework and facilitating interactive predictions allows for flexible, comprehensive, and in-depth analysis of protein functions.

## Methods

### Dataset preprocessing

We collected the amino acid sequences of proteins and their functions from Swiss-Prot (28), the reviewed subset of proteins in UniProtKB (29). The “Function” section in UniProtKB provides a textual description of a protein’s functions. Additionally, the “Keywords” section offers a controlled vocabulary with a hierarchical structure that describes various aspects of protein functions, including activities, locations, interactions, and more. The Swiss-Prot database within UniProtKB, which was manually curated by experts, serves as a high-quality reference for protein functions. The data used in this study was based on the UniProt 2023_02 version, released on May 2nd, 2023^1^. We downloaded the metadata in JSON format and extracted the protein functions by filtering entries where commentType is set to “Function”. We excluded all functions that contain the molecule field, indicating that the function pertains to a subsequence of amino acids after clipping rather than the entire protein sequence. This exclusion is necessary because the protein can serve as a precursor to various chains or peptides. UniProtKB specifies the role of each peptide separately under distinct molecule^2^ entries. As a result, functions for 2,071 proteins were excluded, reducing the total to 523,994 proteins. In our text-based protein function prediction study, we randomly selected 200 proteins to form the test set. For each specific prediction task, 100 proteins were randomly chosen as the test set. The remaining proteins were divided into a training set and a validation set in a 9:1 ratio.

From the training proteins and their associated textual descriptions of functions, we curated the training dataset for ProteinChat (Extended Data Fig. 1). For each training protein *p*, we created a training example represented as a triplet (protein’s amino acid sequence, prompt, answer). The amino acid sequence and the prompt serve as the inputs to ProteinChat, while the answer is the expected output. Specifically, the amino acid sequence of *p* serves as the first element in the triplet, the prompt “Describe the function of this protein” forms the second element, and the textual description of *p*’s function acts as the third element. To enhance ProteinChat’s robustness against linguistic variations, we also employed other semantically equivalent prompts during the training process (22). Additionally, we generated training triplets based on UniProtKB keywords, which are organized into a hierarchy. There are 10 first-level keywords, and we selected 4 that are relevant to protein functions, including molecular functions, binding properties, biological processes, and cellular localization. Furthermore, we chose 31 second-level keywords associated with over 10,000 proteins. These keywords cover 93% of all proteins in Swiss-Prot. Extended Data Table 1 was used to curate training triplets from keywords. For a given protein *p* associated with a keyword *k*, the corresponding prompt *t* for *k* was identified from this table. For example, if the keyword is KW-0808 (“Transferase”), the corresponding prompt is “What type of enzyme is this? Choose one from the following options: hydrolase, oxidoreductase, lyase, transferase, ligase, isomerase, and translocase.” This forms the triplet (*p, t, k*). On average, 2.7 triplets were curated per protein. Extended Data Table 1 presents the number of triplets curated from each keyword. triplets curated from keywords related to molecular function, biological process, and cellular localization cover 67.1%, 35.5%, and 60.8% of all proteins, respectively. The final training dataset for ProteinChat was formed by combining triplets curated from textual descriptions of functions and keywords. Similarly, a validation set of triplets was curated from the validation proteins.

**Extended Data Figure 1.**
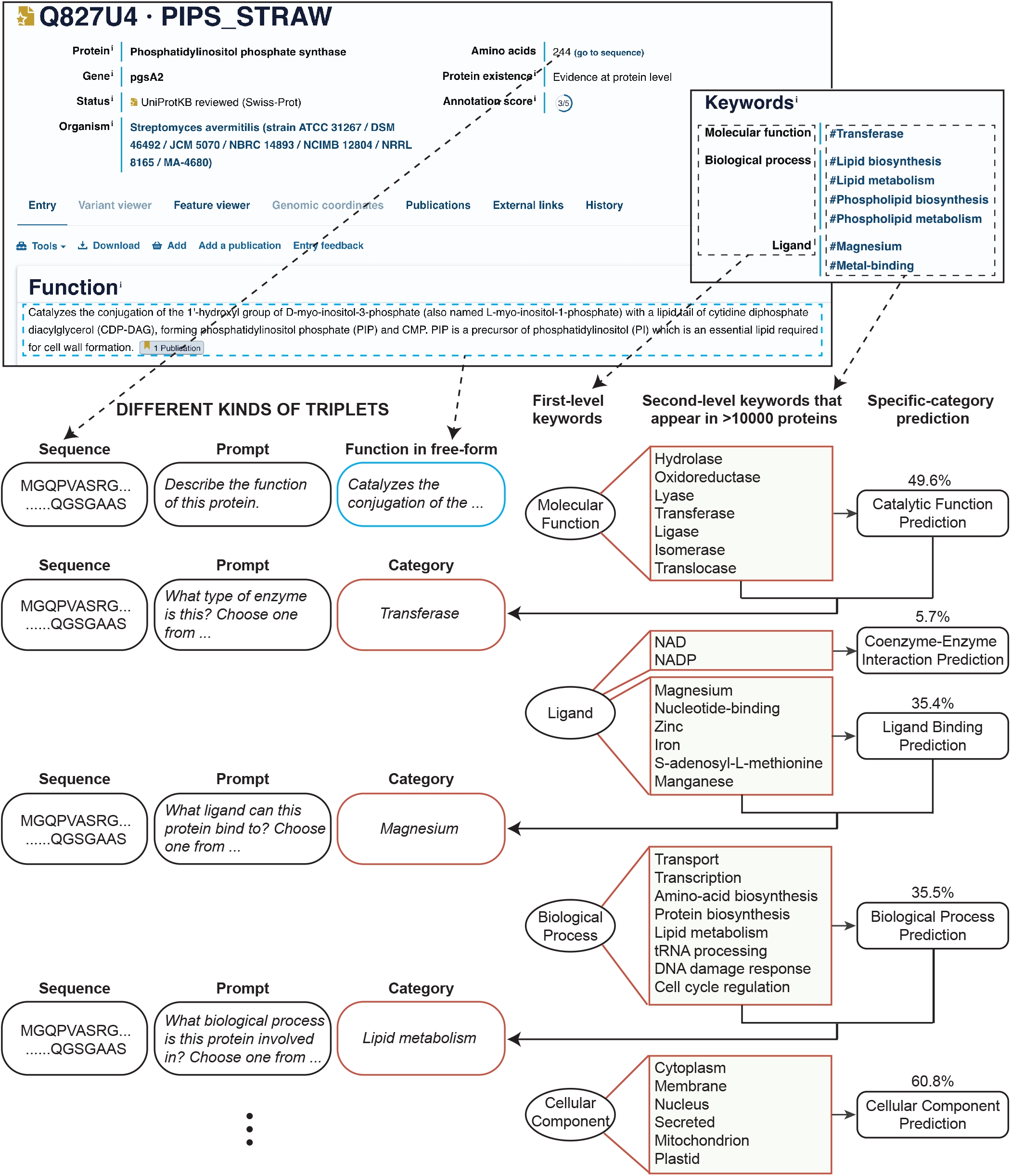
An illustration of the process used to curate (protein sequence, prompt, answer) triplets from the Swiss-Prot database. The percentages represent the percentages of protein entries in Swiss-Prot, whose keywords cover the listed categories.

**Extended Data Table 1.** Prompts linked to keywords and the number of curated triplets for each keyword.

| **Catalytic function** |  |  |  |
| --- | --- | --- | --- |
| **Prompt:** What type of enzyme is this? Choose one from the following options: hydrolase, oxidoreductase, lyase, transferase, ligase, isomerase, and translocase. |  |  |  |
| Function category | Number of triplets | UniProtKB keyword | GO term |
| Transferase | 98540 | KW-0808 | GO:0016740 |
| Hydrolase | 65580 | KW-0378 | GO:0016787 |
| Oxidoreductase | 36864 | KW-0560 | GO:0016491 |
| Ligase | 29379 | KW-0436 | GO:0016874 |
| Lyase | 26546 | KW-0456 | GO:0016829 |
| Isomerase | 16283 | KW-0413 | GO:0016853 |
| Translocase | 14708 | KW-1278 | - |
| **Ligand binding** |  |  |  |
| **Prompt:** What ligand can this protein bind to? Choose one from the following options: magnesium, nucleotide-binding, zinc, iron, S-adenosyl-L-methionine, and manganese. |  |  |  |
| Function category | Number of triplets | UniProtKB keyword | GO term |
| Nucleotide-binding | 101082 | KW-0547 | GO:0000166 |
| Magnesium | 46675 | KW-0460 | - |
| Zinc | 41464 | KW-0862 | - |
| Iron | 29555 | KW-0408 | - |
| S-adenosyl-L-methionine | 17332 | KW-0949 | - |
| Manganese | 12067 | KW-0464 | - |
| **Coenzyme-enzyme interaction** |  |  |  |
| **Prompt:** What coenzyme does this enzyme interact with? Choose one from the following options: nicotinamide adenine dinucleotide (NAD) and nicotinamide adenine dinucleotide phosphate (NADP). |  |  |  |
| Function category | Number of triplets | UniProtKB keyword | GO term |
| Nicotinamide adenine dinucleotide (NAD) | 21502 | KW-0520 | - |
| Nicotinamide adenine dinucleotide phosphate (NADP) | 15102 | KW-0521 | - |
| **Biological process** |  |  |  |
| **Prompt:** What biological process is this protein involved in? Choose one from the following options: molecule transport, DNA to mRNA transcription, amino acid biosynthesis, protein biosynthesis from mRNA molecules, lipid metabolism, tRNA processing, DNA damage response, and cell cycle regulation. |  |  |  |
| Function category | Number of triplets | UniProtKB keyword | GO term |
| Molecule transport | 58648 | KW-0813 | - |
| DNA to mRNA transcription | 32127 | KW-0804 | - |
| Amino acid biosynthesis | 26272 | KW-0028 | GO:0008652 |
| Protein biosynthesis from mRNA molecules | 26063 | KW-0648 | GO:0006412 |
| Lipid metabolism | 16282 | KW-0443 | GO:0006629 |
| tRNA processing | 15380 | KW-0819 | GO:0008033 |
| DNA damage response | 14565 | KW-0227 | GO:0006974 |
| Cell cycle regulation | 14474 | KW-0131 | GO:0007049 |
| **Cellular component** |  |  |  |
| **Prompt:** What is the cellular localization of this protein? Choose one from the following options: cytoplasm, membrane, nucleus, secreted, mitochondrion, and plastid. |  |  |  |
| Function category | Number of triplets | UniProtKB keyword | GO term |
| Cytoplasm | 165882 | KW-0963 | GO:0005737 |
| Membrane | 116756 | KW-0472 | GO:0016020 |
| Nucleus | 41431 | KW-0539 | GO:0005634 |
| Secreted | 32360 | KW-0964 | GO:0005576 |
| Mitochondrion | 17206 | KW-0496 | GO:0005739 |
| Plastid | 15990 | KW-0934 | GO:0009536 |

### ProteinChat model

ProteinChat employs xTrimoPGLM-1B (27) as the protein sequence encoder and Vicuna-13B (25) as the large language model. The xTrimoPGLM-1B model comprises 24 Transformer (37) layers, 32 attention heads, and an embedding dimension of 2048. It was pretrained on the Uniref90 (43) and ColabFoldDB (44) datasets using two strategies: masked language modeling (MLM) (45) and general language modeling (GLM) (46). The MLM strategy enhances xTrimoPGLM-1B’s understanding of protein sequences, while the GLM strategy improves its generative capabilities. Vicuna-13B, fine-tuned from Llama2-13B (21), retains the same architecture as Llama2-13B including 40 Transformer layers, 40 attention heads, and an embedding dimension of 5120. Vicuna-13B was trained by fine-tuning Llama2-13B on a dataset of 70K user-shared dialogues collected from ShareGPT.com.

For an input protein **x**_*p*_, we utilize the pretrained xTrimoPGLM-1B encoder *g* to generate a protein embedding *g*(**x**_*p*_) of size *l ×* 2048, with *l* to be the length of the amino acid sequence. A linear layer (i.e., adaptor) **W** is applied to map these protein embeddings to the LLM input embedding space, resulting in a new embedding **h**_*p*_ = *g*(**x**_*p*_) *×* **W** of size *l ×* 5120. This embedding can be directly input into the LLM to represent the protein. To combine the protein embedding with the textual prompt, we design the LLM Input and Response fields following the conversational format of Vicuna (25):

- (LLM Input) Human: <Protein> `ProteinHere` </Protein> `Prompt` Assistant:
- (LLM Response) `Answer`

As previously mentioned, each training example consists of a (protein, prompt, answer) triplet. We replace the placeholders `Prompt` and `Answer` with the corresponding elements from the triplet. All text in the LLM input, except for `ProteinHere`, is referred to as the *auxiliary prompt*, including the special characters <, >, and /. We denote the tokenized auxiliary prompt as **x**_aux_. Next, we use the LLM to embed **x**_aux_, resulting in the auxiliary prompt embedding **h**_aux_. After obtaining this embedding, we replace `ProteinHere` with the protein embedding **h**_*p*_ generated by the adaptor and feed the entire prompt into the LLM.

The model is trained using a language modeling task, where it learns to generate successive tokens by considering the preceding context. During the training process, the main objective is to optimize the log-likelihood of these tokens. In ProteinChat, only the Answer part is used to compute the loss. By explicitly adding an ending symbol to the answer, the model is also trained to predict where to stop. Specifically, for a target answer **x**_*a*_ of length *l*, we compute the probability of generating **x**_*a*_ by:

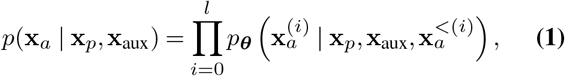

where **x**_*p*_ is the protein sequence and **x**_aux_ is the auxiliary prompt in tokens. **x**_*a*_ is the answer to be trained on. We use 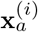 and 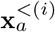 to denote the *i*-th token and all tokens before the *i*-th one. ***θ*** denotes the trainable model parameters.

### Training details of ProteinChat

We used the Adam (47) optimizer with *β*_1_ = 0.9, *β*_2_ = 0.999, and a weight decay of 0.05. We applied a cosine learning rate decay with a peak learning rate of 1*e*-5 and a linear warm-up of 2000 steps. The\ minimum learning rate was 1*e*-6. Due to the high memory consumption required for fine-tuning the encoder and LLM, we utilized a mini-batch size of one per GPU and limited the protein length to a maximum of 600 residues. Notably, 87.1% of the proteins had sequence lengths within this limit. For protein sequences longer than this limit, we truncated the excess length. We used 8 NVIDIA A100 GPUs, with 4 accumulation steps, resulting in an effective batch size of 32. We trained the model for 210K steps. In LoRA, we set the rank to 8, LoRA alpha to 16, and dropout rate to 0.05.

### Evaluation metrics

We employed SimCSE (33) to assess the semantic similarity between the ground truth protein function and the predicted function. SimCSE leverages a contrastive learning framework (48) and utilizes the RoBERTa-base (49) model (denoted by *f*_*θ*_) to generate sentence embeddings. The semantic similarity is quantified by calculating the cosine similarity of these embeddings, with scores ranging from -1 to 1, where higher values signify greater semantic alignment. Specifically, l et *s* a nd *s* ^*′*^ represent the ground truth protein function and the predicted function, respectively. The SimCSE score is computed as:

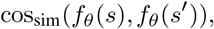

where *f*_*θ*_(*s*) and *f*_*θ*_(*s*^*′*^) are the embeddings of *s* and *s*^*′*^ extracted by the RoBERTa-base model *f*_*θ*_. cos_sim_(*·,·*) denotes the cosine similarity operation.

BLEU (34) is computed using a set of modified n-gram precisions. Specifically,

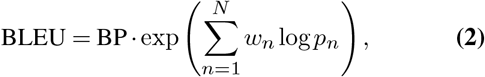

where *p*_*n*_ is the modified precision for n-gram, *w*_*n*_ *>* 0 and 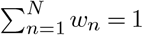. The brevity penalty (BP) is applied to penalize short generated text. Let *c* be the length of the generated text and *r* be the length of the ground truth. BP is computed as follows:

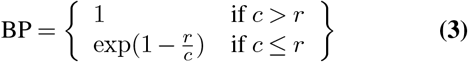

The weighted F1 score is computed by averaging the F1 scores of all categories, taking into account the number of true instances (support) for each category. The macro F1 score is calculated by averaging the F1 scores of all categories without considering their support. The macro F1 score is computed by taking the arithmetic mean (aka unweighted mean) of all the per-category F1 scores, and the weighted F1 score is calculated by taking the mean of all per-category F1 scores while considering each category’s support.

In specific prediction tasks (i.e., classification tasks), both ProteinChat and GPT-4 occasionally produced responses containing multiple answers. For example, a response for biological process prediction might include both molecule transport and amino-acid biosynthesis. Such responses were deemed incorrect, even if they contained the correct answer. We only considered a response correct when it exclusively presented the single correct answer. Additionally, during the evaluation, all texts were standardized to lowercase to avoid the influence of letter casing.

### Experimental details for the GPT-4 baseline

To solicit function predictions from GPT-4 using protein names, we used the following prompt: “You are a biologist specialized in protein functions. Given the name of a protein: [*protein name*], please describe the function of this protein.” When using the amino acid sequence of a protein to solicit function predictions from GPT-4, we used the following prompt: “Given the sequence of a protein: [*a string of amino acid letters such as MARYFRRRKFCRFTAEGVQEIDYKDIATLKNYITESGKIVPSRITGTRAKYQRQLARAIKRARYLSLLPYTDRHQ*], please describe the function of this protein.”

### Experimental details for specific prediction tasks

Predicting enzyme catalytic functions involves determining which of the seven categories of chemical reactions a given enzyme can catalyze. These categories include hydrolase, oxidoreductase, lyase, transferase, ligase, isomerase, and translocase. The prompt for this prediction task was “What type of enzyme is this? Choose from [*the list of categories above*]”. Similarly, predicting ligand binding entails identifying the specific ligand a protein can bind to, while predicting coenzyme-enzyme interactions focuses on determining which coenzyme interacts with a given enzyme. The prompts for these tasks are outlined in Extended Data Table 1. In the biological process prediction task, the goal is to predict the biological processes in which a protein is involved, including molecule transport, DNA to mRNA transcription, amino acid biosynthesis, protein biosynthesis from mRNA molecules, lipid metabolism, tRNA processing, DNA damage response, and cell cycle regulation. Cellular component prediction involves determining the cellular localization of proteins (32). While cellular localization does not directly define protein functions, it is often intrinsically linked to the roles proteins play within the cell. For example, proteins involved in energy production, such as those in the electron transport chain, are typically located within the mitochondria. We evaluated ProteinChat’s ability in identifying proteins’ cellular localization from six categories: cytoplasm, membrane, nucleus, secreted, mitochondrion, and plastid, using the following prompt: “What is the cellular localization of this protein? Choose from [*a list of the six categories*]”.

For each of these specific prediction tasks, we developed a specialized classifier. Each classifier includes a protein encoder based on the pretrained xTrimoPGLM-1B and a classification head based on a multi-layer perceptron. Given the amino acid sequence of a protein, the protein encoder extracts representations for each amino acid. These representations are then averaged into a single vector, which is subsequently fed into the classification head to predict the class label. The classification head is a Multilayer Perceptron (MLP) with two layers. For all classification tasks, the first layer of the MLP contains 128 hidden units. The second layer’s number of hidden units corresponds to the number of categories specific to the task. For each classifier, we trained two variants: 1) keeping the pretrained protein encoder fixed and only training the classification head (referred to as Classifier 1), and 2) training both the protein encoder and the classification head (referred to as Classifier 2). The weights of the MLP were initialized using the Kaiming initialization method. We used the same learning rate and optimizer as in the ProteinChat training configurations. The batch size was set to 32, and a checkpoint was saved every 2500 iterations. The checkpoint with the best performance on 300 randomly selected validation examples was then chosen. For each task, there were 100 test proteins. The training data for the specialized classifiers was curated from the UniProtKB database. The number of training examples for the classifiers in the tasks of predicting catalytic functions, ligand binding, coenzyme-enzyme interactions, biological processes, and cellular components were 277548, 198215, 31672, 340276, and 198661 respectively.

The two Gene Ontology (GO) classifiers - DeepGO-Plus (35) and NetGO 3.0 (36) - utilize online web services to predict GO terms with rankings. A prediction is considered correct if the ground truth GO term holds the highest rank among all possible answers for the given question.

### Use GPT-4 to assess ProteinChat’s text-based predictions of protein functions

GPT-4 has demonstrated effectiveness in assessing the quality of text generated by large language models. We utilized GPT-4 to assess the accuracy of ProteinChat’s text-based function predictions by comparing them with the ground truth descriptions. The specific prompt provided to GPT-4 for this evaluation is: “You are a biologist specialized in protein functions. Please compare the predicted function ‘[*predicted function*]’ with the ground truth function ‘[*ground truth function*]’. Then give a score based on the following rubric. Assign a score of 2 if the predicted function is an exact match to the ground-truth function, or it is a subset of the ground-truth function. Assign a score of 1 if some aspects of the predicted function align with the ground truth but other aspects conflict with it. Assign a score of 0 if the predicted function does not align with the ground truth at all.” The evaluation rubric mirrored that of human expert assessments, consisting of scores 2, 1, and 0. GPT-4 assigned an average score of 1.36 to ProteinChat’s predictions for the 200 test proteins (Extended Data Fig. 2a). In contrast, GPT-4’s own generated predictions received a significantly lower average score of 0.17. The correlation between the evaluation results of human experts and GPT-4 was 0.72, indicating a strong agreement.

**Extended Data Figure 2.**
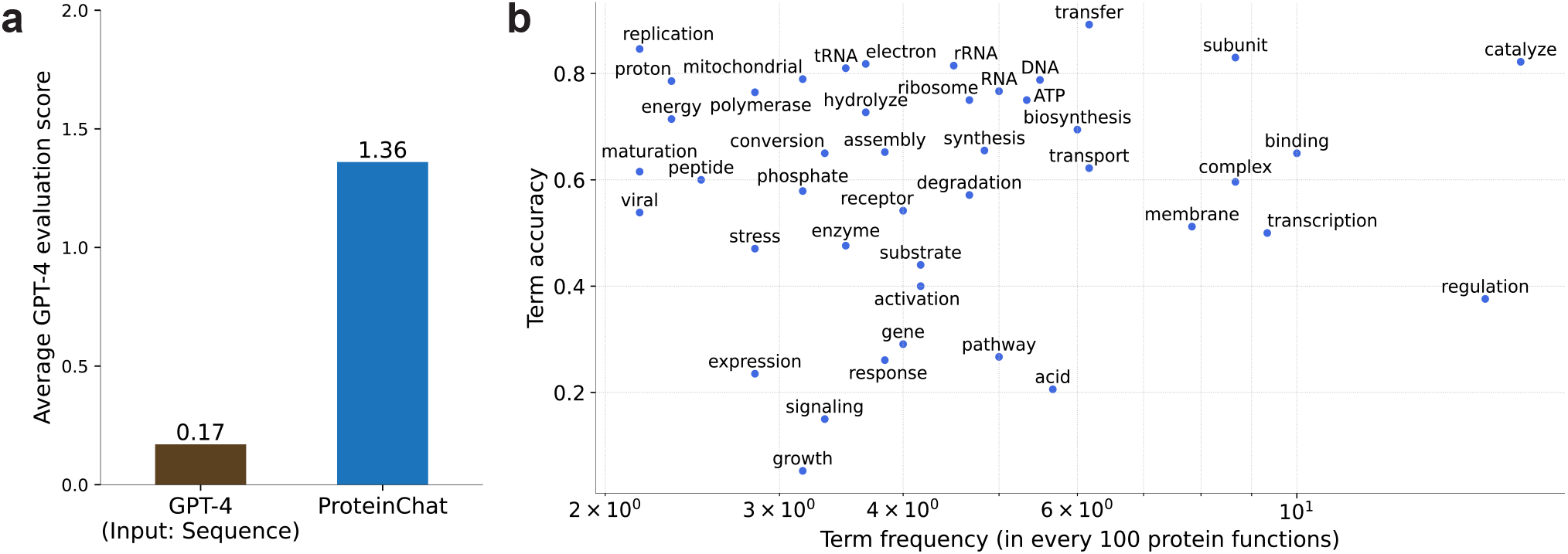
**a**, GPT-4 evaluation scores for ProteinChat, compared to GPT-4 predictions using protein sequences as input. **b**, ProteinChat’s prediction accuracy for biological terms across varying frequencies.

### ProteinChat accurately predicts biological terms

To further evaluate the correctness of the text-based protein functions predicted by ProteinChat, we introduced an additional evaluation metric called Biological Term Accuracy. We collected a set of biological terms and assessed the accuracy for each term *t* as follows: For each test protein, if *t* is either present or absent in both the protein’s ground truth function description and the function predicted by ProteinChat, then the prediction is considered correct. Otherwise, it is considered incorrect. The accuracy for *t* is defined as the ratio of the number of correct predictions to the total number of test proteins. To collect a vocabulary of biological terms, we utilized SciSpacy (50), a Python library tailored for biomedical and scientific text processing, to extract biological terms from 600 randomly sampled ground truth function descriptions. From these extracted terms, we selected the 43 most frequently occurring terms. Extended Data Fig. 2b shows the accuracy of these terms versus their frequency on a logarithmic scale. ProteinChat achieved high accuracy on the majority of these terms, demonstrating its capability to capture key biological information in its predictions.

### Proteins with identical functions are located close to each other in the representation space of ProteinChat

To better understand how ProteinChat predicts protein functions, we visualized its learned protein representations in a 2D space using t-SNE (51). For each input protein’s amino acid sequence, we utilized the trained xTrimoPGLM (27) protein encoder and the trained adaptor in ProteinChat to extract a representation vector for each amino acid. We then computed the overall representation of the entire protein by averaging the representations of all the amino acids. We projected the protein representation vectors into a 2D space using t-SNE (51) for visualization. Extended Data Fig. 3 presents a visualization of all *n* = 20, 426 human proteins from the Swiss-Prot dataset. Each dot in the figure represents a protein. In Extended Data Fig. 3a, we have highlighted proteins with ground truth labels for three cellular localizations: nucleus (*n* = 5, 617), secreted (*n* = 2, 113), and mitochondrion (*n* = 1, 309). As observed, proteins with the same cellular localization are clustered together in the representation space. Similar patterns can be observed in Extended Data Fig. 3b-d. This demonstrates ProteinChat’s ability in grouping functionally similar proteins together, thereby enhancing the accuracy of function predictions.

### Impact of hyperparameters

We investigated how the hyperparameters used during text generation in ProteinChat affect the quality of the generated text. Extended Data Fig. 4 show the average BLEU-1 (higher is better) and perplexity (PPL, lower is better) scores when varying beam search depth (the number of top results maintained during the search for the best responses) and temperature (the likelihood of sampling low-probability tokens). Our findings show that the performance remains relatively stable across different beam search depth values. On the other hand, we observed that a higher temperature slightly decreases generation performance. This is likely because higher temperatures encourage more diverse and less predictable token selection, which can lead to the generation of less coherent and grammatically incorrect sentences.

**Extended Data Figure 3.**
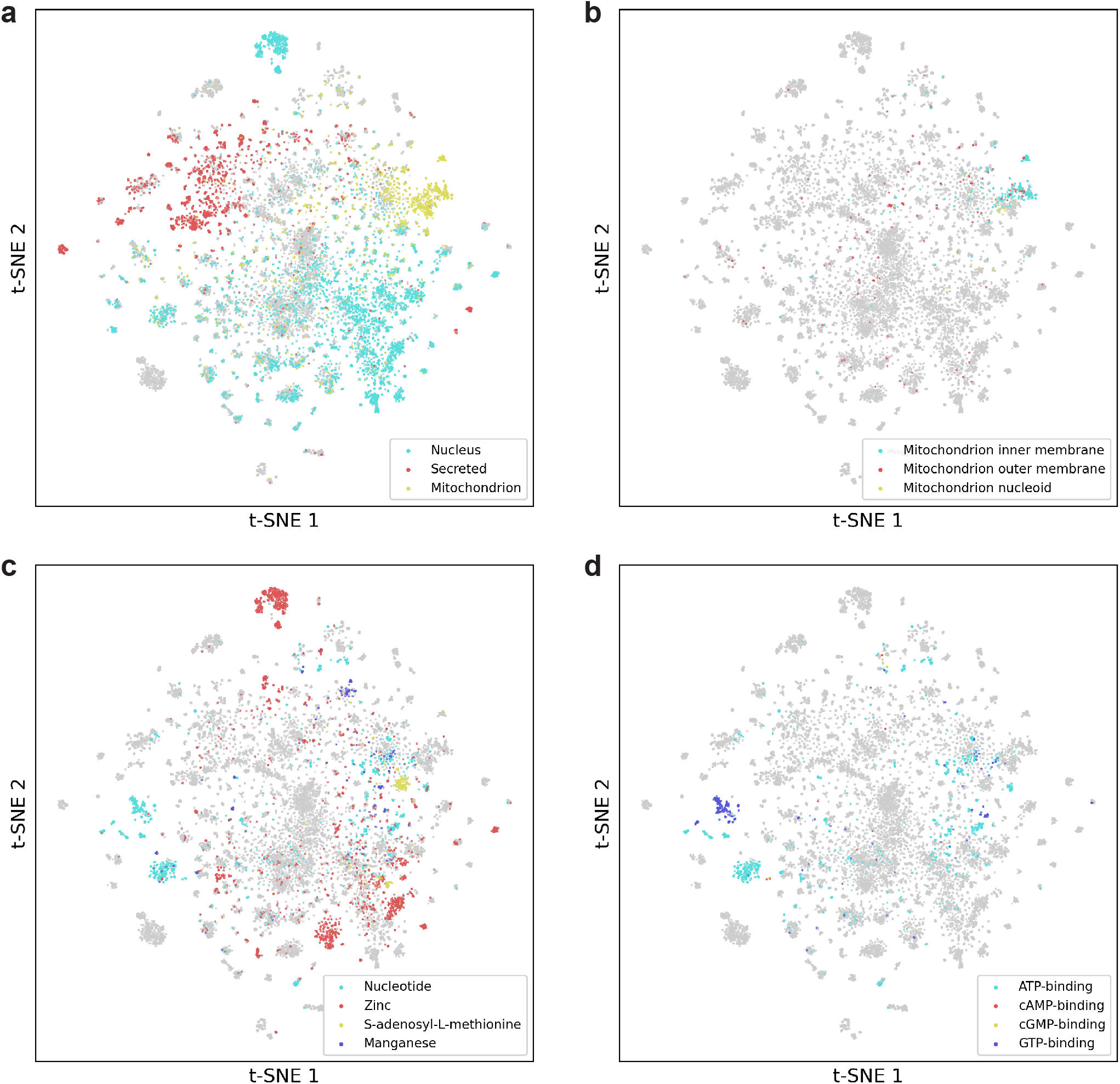
t-SNE visualization of protein representations extracted by the protein encoder and adaptor of ProteinChat. **a**, Proteins located in three cellular locations, including nucleus, secreted, and mitochondrion, are highlighted. **b**, Proteins located in three mitochondrial components inner membrane, outer membrane, and nucleoid - are highlighted. **c**, Proteins that bind with four ligands - nucleotide, zinc, S-adenosyl-L-methionine, and manganese - are highlighted. **d**, Proteins binding with ATP, cAMP, cGMP, and GTP are highlighted.

### Related work

To better analyze, annotate, and predict protein functions, significant research has been conducted in recent years. The Critical Assessment of Function Annotation (CAFA) competition (7) is designed to develop machine learning models for predicting the Gene Ontology (GO) categories associated with protein functions. As of 2023, this competition has been held five times, yielding diverse solutions such as comparing unsolved sequences with known proteins, integrating multiple data sources, and applying machine learning algorithms with insights into biological processes to decipher protein functions. Notable work has focused on predicting GO functions, including DeepGOPlus (18, 35) and NetGO 3.0 (36). These methods typically train separate models for each sub-ontology in GO, which encompasses molecular function ontology (MFO), biological process ontology (BPO), and cellular component ontology (CCO). Recent deep learning methods have demonstrated great efficacy in predicting specific protein functions. These include Graph Neural Networks (13), diffusion models (3), transfer learning (52), and contrastive learning (17). These methods focus on predicting protein functions represented as discrete categories, but they are unable to predict functions described in free-form text, which typically contains more detailed information than category labels.

**Extended Data Figure 4.**
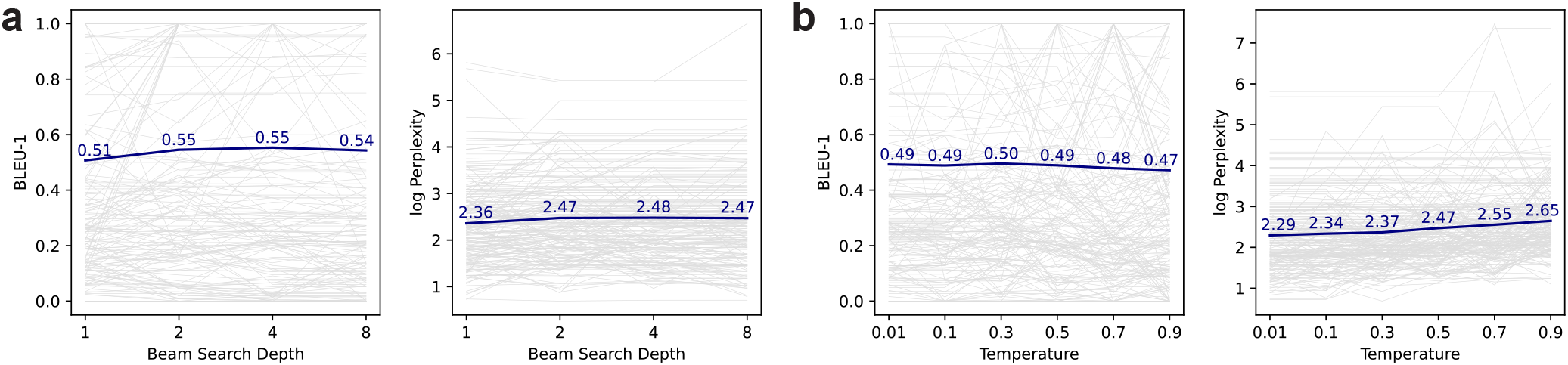
BLEU-1 and perplexity scores of text-based protein functions predicted by ProteinChat, evaluated under different beam search depths (**a**) and temperatures (**b**).

Multi-modal learning, particularly in image-text applications, has seen significant advancements recently. The CLIP model (53) employs contrastive learning to align image and text embeddings effectively. The BLIP-2 framework (54) integrates images and text prompts to generate relevant responses using large language models. Building on BLIP-2, MiniGPT-4 (22) enhances performance by incorporating the more powerful Llama-2 model. Additionally, LLaVA (55) combines a vision encoder with a large language model for various visual-textual tasks, including scientific question answering. In the scientific domain, multi-modal learning has gained increasing attention. MoleculeSTM (56) utilizes contrastive learning to simultaneously learn representations for chemical structures and textual descriptions of molecules. ProtST (57) employs contrastive learning and multi-modal mask prediction to align protein sequences with their textual descriptions, enabling zero-shot classification and text-protein retrieval. In contrast to ProtST, ProteinChat offers free-form protein function prediction, a feature not available in ProtST. Additionally, MultiVI (58) is a deep generative model that integrates multi-modal single-cell datasets, facilitating the joint analysis of chromatin accessibility and gene expression measurements.

## Data availability

All data used in this study are available at https://drive.google.com/file/d/1n5Ant3S5QE0Yx-DznRa3lannFanc1WB7/view?usp=sharing.

## Code availability

The source code of this work is available at https://github.com/mignonjia/ProteinChat. We use ESM-1B (9) instead of xTrimoPGLM as the protein encoder in this GitHub repository because xTrimoPGLM is currently not publicly available.

---

**Extended Data Table 2.** Rubric for human expert assessment of predicted protein functions.

| Summary | Criteria | Score |
| --- | --- | --- |
| Correct | The predicted function satisfies one of the following criteria:<br>1) It is an exact match to the ground-truth function.<br>2) It is a subset of the ground-truth function.<br>3) It contains additional, accurate information beyond the ground-truth function.<br>4) It does not directly align with the ground-truth function but represents another correct function for the protein. This can be verified through domain knowledge or by checking the publication associated with this protein on UniProtKB. | 2 |
| Partially Correct | While some aspects of the predicted function align with the ground truth, other aspects conflict with it. | 1 |
| Incorrect | The predicted function meets one of the following criteria:<br>1) It is entirely inaccurate.<br>2) It is irrelevant to the question. | 0 |
| Ambiguous | It lacks information to make a comparison between the predicted function and the ground truth function. | - |

**Extended Data Figure 5.**
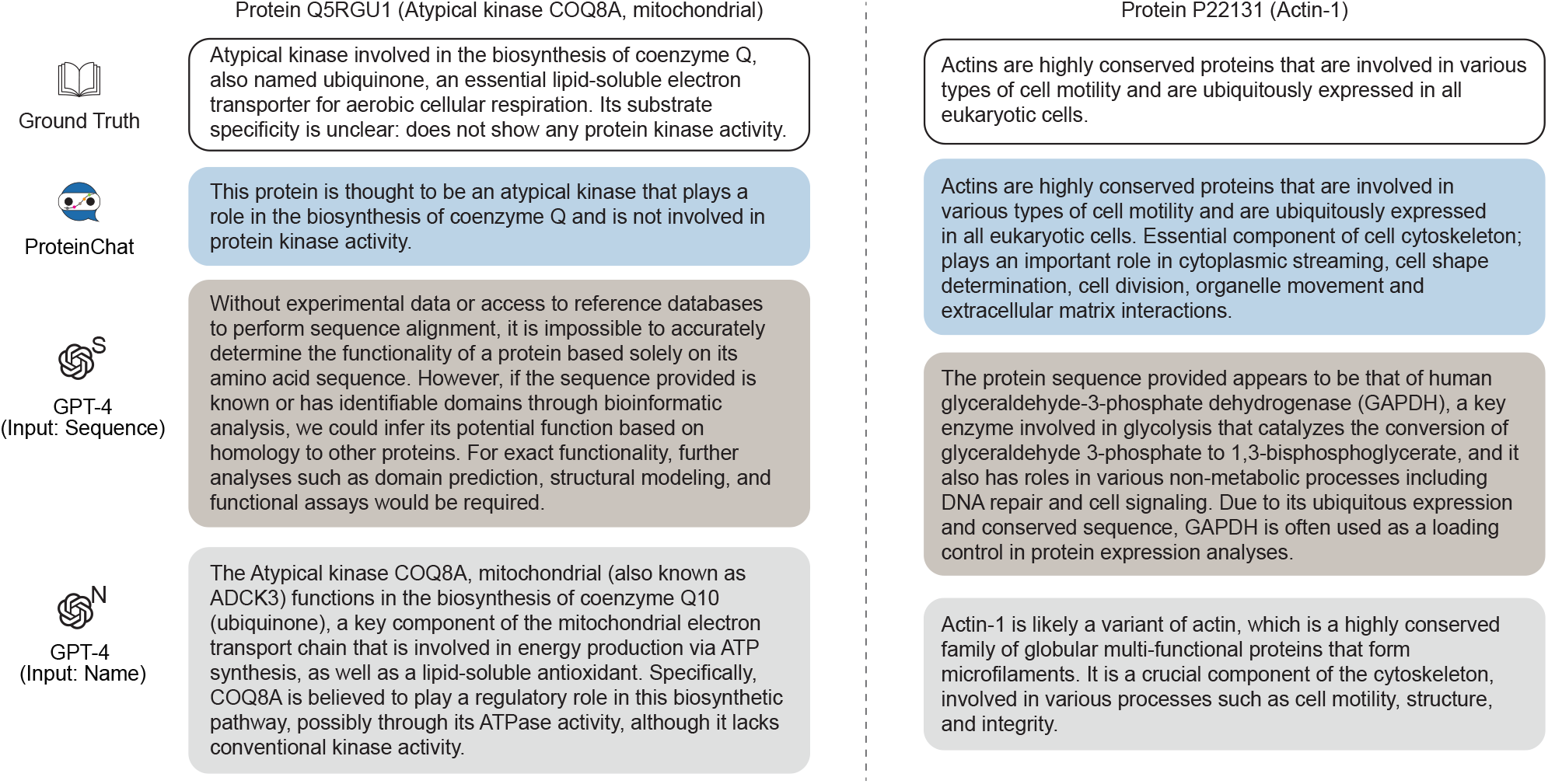
Comparison of predictions generated by ProteinChat and GPT-4 using amino acid sequences or protein names as inputs for two additional randomly selected test proteins.

## Footnotes

1 https://www.uniprot.org/release-notes/2023-05-03-release

2 https://www.uniprot.org/help/function

## Notes

### Competing Interest Statement

The authors have declared no competing interest.

